# Yn-situ: a robust single RNA molecule *in situ* detection method

**DOI:** 10.1101/2021.10.20.465061

**Authors:** Yunming Wu, Wenjing Xu, Limei Ma, Zulin Yu, Yongfu Wang, C. Ron Yu

**Author notes:** Current address: Department of Biology, Howard Hughes Medical Institute, Stanford University, Stanford, CA 94305, USA.

## Abstract

We describe a cost-effective, highly sensitive, and quantitative method for *in situ* detection of single RNA molecules in tissue sections. This method, dubbed *Yn situ*, standing for Y-branched probe *in situ* hybridization, uses a single-strand DNA preamplifier with multiple initiation sites that trigger hybridization chain reaction (HCR) to detect polynucleotide. We characterized the performance of this method and compared it to other approaches in the postnatal mouse olfactory epithelia. We find that the *Yn situ* method, in conjunction with an improved fixation step, is sensitive enough to allow detection of single molecules using a single pair of probes targeting a short nucleotide sequence. A set of 5-probes can produce quantitative results with smaller puncta and higher signal-to-noise ratio than the 20-probe sets commonly required for HCR and RNA-Scope. We show that the high sensitivity and wide dynamic range allow quantification of genes expressed at different levels in the olfactory sensory neurons. We describe key steps of this method to enable broad utility by individual laboratories.

## Introduction

Detecting nucleotide acid using *in situ* hybridization has been an important methodology in biological sciences since it was first invented in 1968 and remains the gold standard of RNA detection in the cell (Gall and Pardue, 1969). Over the decades, various techniques have been developed to improve sensitivity, specificity, resolution, quantification, and to simultaneously detect multiple targets (Chen et al., 2015; Choi et al., 2010; Ishii et al., 2004; Shah et al., 2016; Wang et al., 2012; Wang et al., 2018). The primary challenge to *in situ* detection of polynucleotides is multifold. First, RNA is unstable in biological samples because of the ubiquitous presence of RNases. Degraded RNAs can lead to diffusive signals that increase background noise. Second, hybridization conditions may vary depending on the length and composition of the probes. The length of the target also limits its detectability. Small RNA and short open reading frame transcripts have fewer specific targeting sequences. Third, there usually is a trade-off between sensitivity and specificity. For example, high intensity signal from methods based on catalyzed reporter deposition (CARD) is usually accompanied by high background noise (Ishii et al., 2004). Detection using directly labeled nucleotide acids has high specificity, but the signal is relatively weak (Trcek et al., 2012). To improve probe stability and specificity, short DNA oligos, especially split probes have been adopted in both RNA-Scope and hybridization chain rection (HCR) protocols (Pizzorusso et al.; Wang et al., 2012). These new methods also employ multiple probes for the same target to improve sensitivity (Trcek et al., 2012), enhance signals through hybridization-based amplification (Zhang et al., 2018), or both (Choi et al., 2018; Wang et al., 2012). These significant improvements allowed the quantification of single molecules using fluorescent signals. However, the high number of specific probes required by these methods incur high costs and can be limiting because only long polynucleotide molecules can provide sufficient target sites. Here we present a new, cost-effective method of *in situ* hybridization that requires significantly fewer probes while achieving equal or superior sensitivity, specificity, spatial resolution, and dynamic range when compared to other contemporary methods.

## Results

### Design of *Yn situ*

The design of *Yn situ* and general procedures are illustrated in Figure 1. The method has improved upon previous approaches in three aspects. First, we have adopted a preamplifier design to allow a single probe to amplify signal multi-fold. In HCR (v3.0), multiple pairs of target probes were used to increase sensitivity. Each pair of target probes hybridize specifically to their binding sites on the target mRNA. The two probes are adjacent to each other such that the un-hybridized portion of the oligos forms an initiator to enable cooperative initiation of HCR reaction. We have modified this design by redesigning the probe pairs such that the un-hybridized sequence is targeted by the preamplifier. The preamplifier probe, when hybridized to the target-specific probes, forms a Y-shaped structure, hence the namesake of this method (Figures 1A). Each preamplifier carried 20 initiator repeats, which can simultaneously trigger 20 HCR reactions (Figures 1A and 1C). This design maintains the use of short oligo sequences (52 nt) and paired probes as in HCR to improve specificity. On the other hand, the use of a preamplifier significantly increases sensitivity while avoiding the requirement of many probe pairs to generate significant signals. Second, we have designed a strategy to generate pre-amplifier that can be readily made with basic molecular biology (Figure 1B-1D). Third, we have adopted a chemical modification of cellular RNAs to reduce RNA degradation and effectively improve staining quality. Previous studies have identified carbodiimide fixatives that effectively crosslink the phosphate group of the cellular RNA with amine groups from the proteins (Pena et al., 2009; Sylwestrak et al., 2016). We thus developed a protocol to irreversibly immobilizing RNA molecules by crosslinking them to formaldehyde-fixed proteins using 1-ethyl-3-(3-dimethylaminopropyl) carbodiimide (EDC).

**Figure 1.**
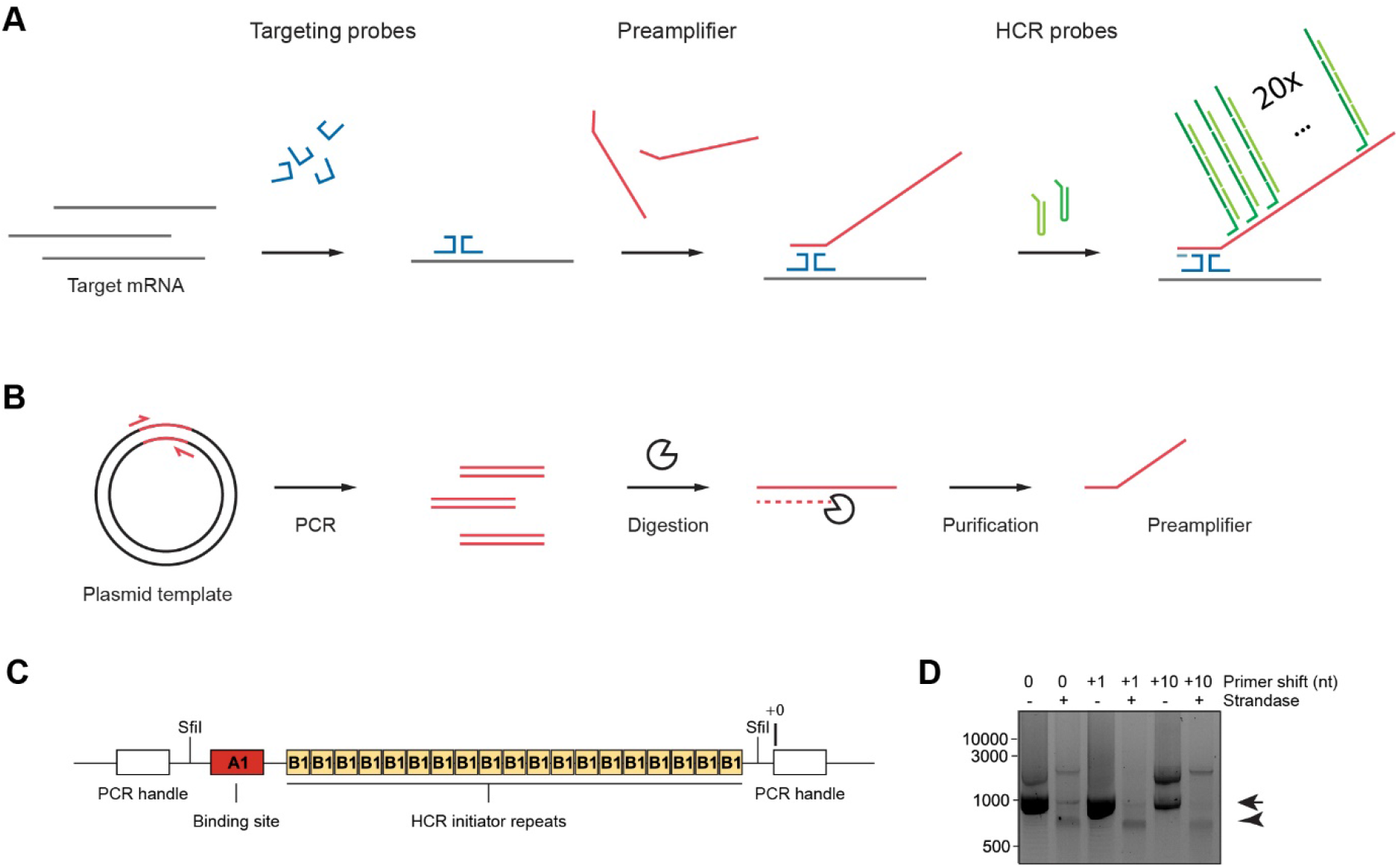
Schematic illustration of Yn situ hybridization. **A**. Steps involved in the hybridization processes. The target RNA is fixed to the cellular proteins by covalent bonds to prevent degradation. A pair of targeting probes (dark blue) recognizes a consecutive 52 nt sequence of the target. A pre-amplifier probe (red) recognizes the tail sequences only when the two targeting probes are aligned next to each other with head-to-head orientation. Each pre-amplifier probe carries 20 HCR initiation sites. Upon incubation with fluorescently labeled metastable HCR hairpins (green and dark green), the HCR initiation sites trigger enzyme independent amplification through HCR resulting in bright fluorescent signals. **B**. process of synthesizing the preamplifier. PCR amplicons are digested with strandase to release single-stranded preamplifier probes. **C.** Schematic illustration of the pre-amplifier probe. The pre-amplifier probe sequence contains a targeting probe binding site (A1) and 20 HCR initiator sequences (B1). The sequence is flanked by two SfiI sites for verification purpose. The sequence including the SfiI sites is flanked by two primer binding sites that allow for exponential amplification using PCR. **D**. Pre-amplifier probe synthesis with different PCR primers. The expected PCR product and ssDNA probe are indicated by arrow and arrowheads, respectively.

### Design and synthesis of the preamplifier probe

The preamplifier contains a binding site to the paired probes and 20 repeats of HCR initiators (Figure 1C). Although the design is simple, it presents a challenge to generate the oligos. Because each preamplifier is approximately 1kb long and contains repetitive sequences that serve as initiation site, it is difficult to synthesize directly. We designed a plasmid that contained the double stranded version of the preamplifier sequence (Figure 1B). The double stranded preamplifier sequence on the plasmid was flanked by sequences recognized by the restriction enzyme SfiI and a pair of PCR handles that are used to amplify the fragment. SfiI digestion could be used to verify the total length of the preamplifier. Double stranded amplicons are generated using asymmetric PCR primers (one with 5’ phosphate, one without). The 5` phosphate on the reverse primer allows the strandase to digest the antisense strand and produce the single-stranded preamplifier.

To determine the optimal condition and kit for PCR amplification of the preamplifier, we first tested 5 commercial PCR polymerases: PrimeSTAR HS DNA polymerase, KAPA HiFi DNA polymerase, Q5 high-fidelity DNA polymerase, LongAmp Taq DNA polymerase, and GoTaq long PCR master mix. We found KAPA HiFi and Q5 polymerase generated nonspecific products. PrimeSTAR and GoTaq long PCR mix generated the desired band at a low yield. LongAmp polymerase generated the desired band with the highest yield (Figure S1A). We chose LongAmp for further optimization. We tested various PCR parameters including annealing temperature, primer concentration, template concentration, and the choice between two-step or three-step PCR for LongAmp. We found the desired product can be generated at almost any annealing temperature tested except 72°C (Figures S1B and S1C). The three-step PCR generated some smear bands below the target band (Figure S1C). This did not affect the experiment since the band was further purified by gel extraction. The optimal primer concentration was 0.5 μM. The optimal template concentration was 0.05 ng/μL among the conditions tested (Figure S1D). Finally, we found that the strandase activity was influenced by the sequences at the priming site. This influence was not clearly understood. We, therefore, empirically tested a series of reverse primers to determine the optimal site for strandase digestion. We identified that priming at +1 and +40 nt away from the initial PCR handle site (+0) produced the most complete digestion (Figure S1E).

Note that the PCR amplification process produced preamplifiers containing the PCR handle sequences. This may increase background noise if these parts of the probe bind to complementary sequences in the cell. To reduce this background, short oligos corresponding to the PCR handle sequences were used at 10 times the preamplifier concentration as a blocking reagent in further experiments.

### Determining the optimal condition for hybridization

We next performed a series of tests to determine the optimal experiment condition for *Yn situ* using probes against olfactory marker protein (Omp), a marker gene expressed by the mature olfactory sensory neuron (mOSN) (Figure 2). Since the probe design and hybridization condition was the same as HCR, we used the same concentration of target probes. We found that 5 pairs of target probes of *Yn situ* produced highly specific and stronger signals than the 20-probe pairs for HCR (Figure 2 A and B). Using the 5 probe pairs we varied the concentration of the preamplifier. Signals can be detected with preamplifier concentration as low as 0.002 ng/μL but lowering preamplifier concentrations significantly reduced the number of spots detected (Figure 2C-D). There was no signal produced when the antisense sequence of the preamplifier was used, indicating that the signals produced by *Yn situ* were highly specific (Figure 2E).

**Figure 2.**
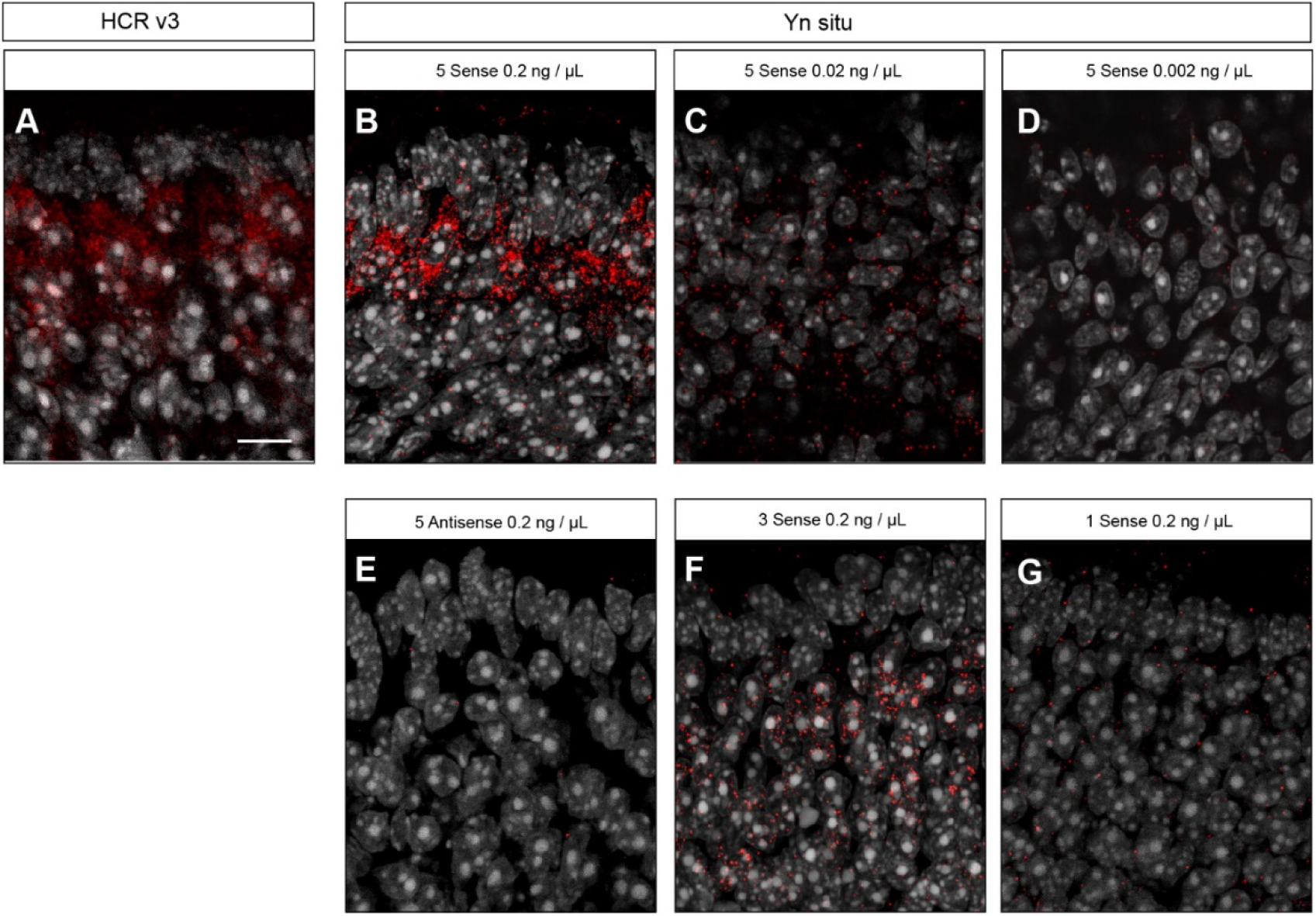
Optimal condition for *Yn situ* hybridization. **A**. A representative image showing the spatial localization of Omp mRNA detected in the olfactory epithelium (postnatal day 0-3) by 3^rd^ generation HCR in situ hybridization. **B-D**. Representative images showing the spatial localization of Omp mRNA detected in the olfactory epithelium by *Yn situ* hybridization using 5 probe pairs with different preamplifier concentrations. **E**. A representative image showing the *Yn situ* hybridization signals using antisense preamplifier probe as a negative control. **F-G**. Detection of Omp signal using 3 pairs (F) and a single pair (G) of probes,

We also tried to determine the minimal pairs of targeting probes required for producing visible signals. Strong signals were detected using 3 pairs of probes (Figure 2F). Even one pair of targeting probes produced signals for Omp, but the signal was weak and not suitable for quantitative studies (Figure 2G). The best results were achieved at 0.2 ng/μL preamplifier, room temperature for HCR with 60 nM of each hairpin (Figure 2B). Hairpin concentration lower than 60 nM did not produce any visible signal (data not shown).

### Characteristics of the *Yn situ* signals

We compared the signals generated by *Yn situ* with those by conventional CARD reactions and the contemporary methods (Figure 3). As recommended by manufacturers, 20 probe pairs were used for RNA-Scope and 3^rd^ generation HCR *in situ*. We conducted super-resolution microscopy using Leica Hyvolution, which was a deconvolution method based on the point spread function (PSF) to allow high-speed multicolor imaging with a resolution down to 140 nm (Borlinghaus and Kappel). Unlike the diffuse signals developed using alkaline phosphatase (AP, Figure 3A) and horseradish peroxidase (HRP, Figure 4A), the fluorescent signals generated by *Yn situ* are small puncta as those found with HCR and RNA-Scope (Figure 3A). Moreover, unlike non-specific signals from AP or HRP reaction, few signal puncta were observed for *Yn situ* in the cells that did not express the target gene. We calculated the signal to noise ratio (SNR) of the four methods (Figure 3B). Because the experimental conditions are different for each method, for comparison, we normalized the signal intensity and used the variance of background signals to calculate SNR. We found AP generated signals had highest relative background noise. RNA-Scope and *Yn situ* had the narrowest distributions of background noise signal. They also have similar distributions of signals detected in the puncta, which were tighter than those generated by AP and HCR. *Yn situ* had the highest signal-to-noise ratio even with less pairs of targeting probes.

**Figure 3.**
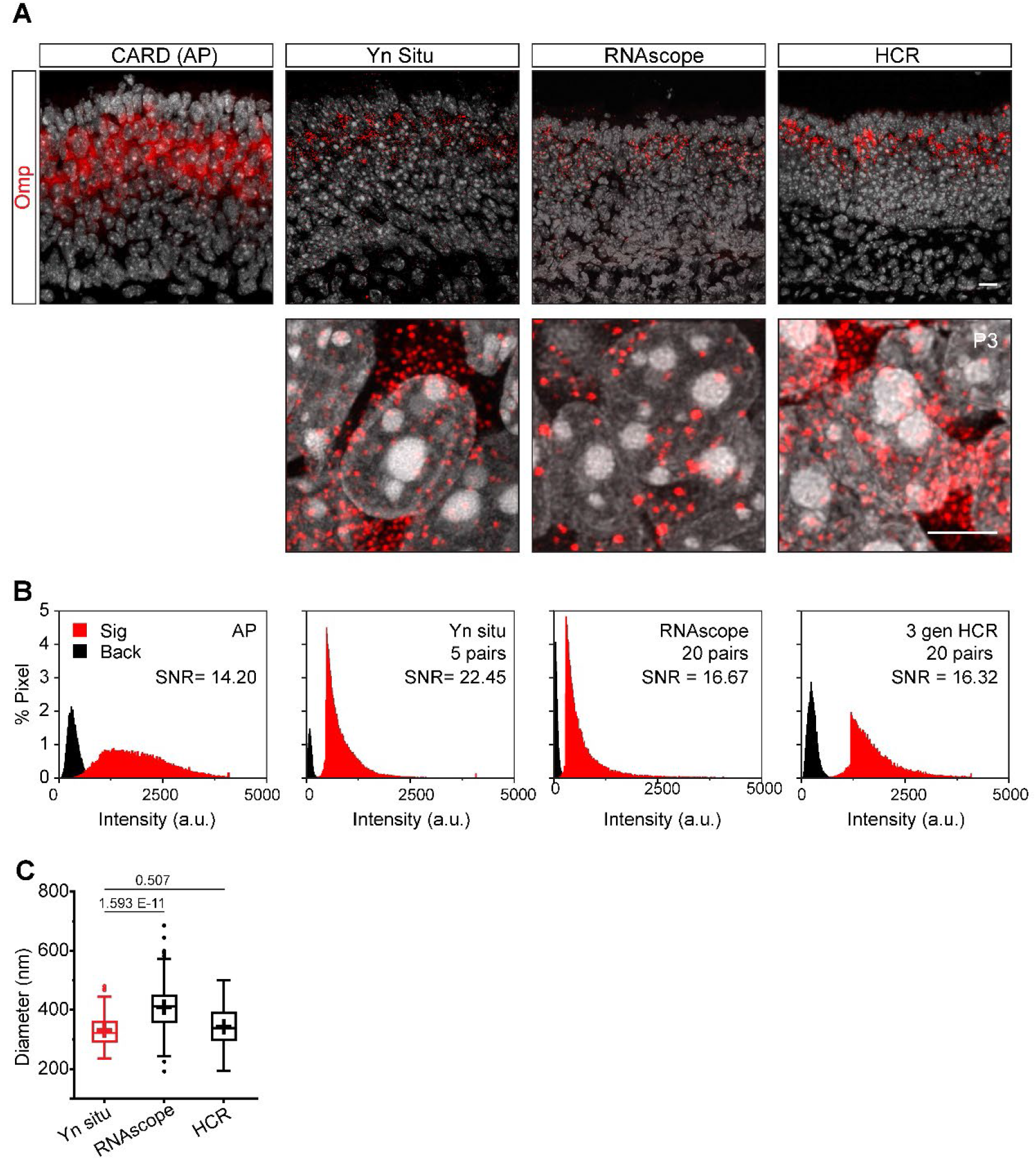
Characterization of the *Yn situ* signals. **A**. Representative images showing the spatial localization of Omp mRNA detected in the olfactory epithelium by conventional FISH using AP (left panel), *Yn situ*, RNAscope and HCR. High magnification pictures are shown in the lower panels. Scale bar, 10 μm. **B**. Histogram of signal strength from pixels in the puncta (red) and background (black). Signal-to-noise ratio (SNR) of the methods are calculated accordingly. The pixel intensity was normalized between 0 – 4095 for comparison between from different experiments. **C**. Box plot showing the puncta sizes measure for the three methods examined. P values are show above the boxes.

**Figure 4.**
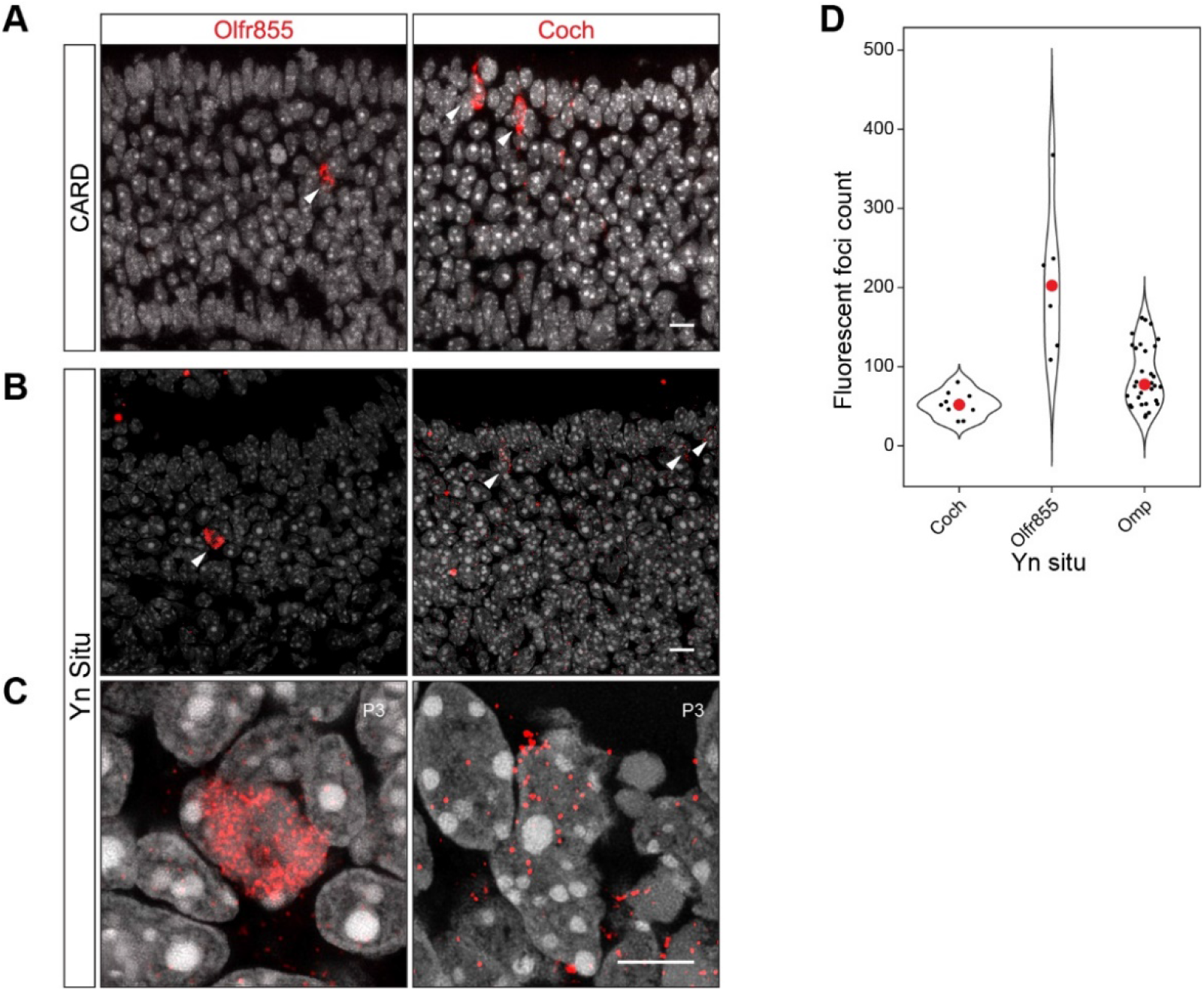
Dynamic range of the *Yn situ* signals. **A.**Representative images showing signals from traditional RNA in situ hybridization in detecting Omp and cochlin (Coch) respectively in the olfactory epithelium using HRP. The signal strengths in individual cells appear similar for the two genes. **B-C**. Representative images showing the *Yn situ* signal for the same genes. Individual signal puncta are clearly visible in the high-resolution images (C). Scale bar, 10 μm. **D**. Quantification of the number of signal puncta in each cell for three genes.

We also determined size of the fluorescent foci for the three contemporary methods (Figure 3C). The puncta sizes for *Yn situ* was significantly smaller than RNA-Scope (Wang et al., 2012). It was also smaller than the puncta from HCR V3.0, although the difference was not statistically significant.

#### Dynamic Range

We have performed *in situ* hybridization experiments against several genes that had variable expression levels. In the olfactory epithelium, all mature OSNs express OMP at the intermediate level. Individual olfactory receptor (OR) genes were expressed at high levels by a very few neurons. A small population of cells expressed the gene cochlin at a moderate level. In CARD experiments, signals amplified from enzymatic reactions often obscured the quantity of RNA in these cells. The signal in neurons expressing an OR appeared similar to those expressing cochlin (Figure 4A). *Yn situ*, on the other hand, allowed a wide range of expression levels of RNA transcription to be quantified (Figure 4B-D). Cochlin signals were comparatively lower than that of OMP. For Olfr855, we detected nearly 400 puncta in a single cell without the signals being saturated. This result demonstrated a high dynamic range of detection by *Yn situ*.

## Discussion

In this study, we have conducted a proof-of-principle study of the *Yn situ* method using perinatal olfactory epithelium sections. Although it has not be tested against other tissues or at different developmental time point, the data we have collected demonstrate that the method can produce high quality and quantitative detection of RNA species. Further optimization likely will enhance the utility of this method. *Yn situ* offers five advantages over current approaches.

First, the *Yn situ* signal is highly specific. The SNR is highest among all of the methods tested even though it uses the least pairs of probes. The signal puncta are spherical. With a diameter of ~300 nm (5 pairs of primary probes), they are also smaller than other methods. The signal is bright and makes it visible directly under the microscope.

Second, the small puncta size enables digital quantification of the RNA transcripts even for highly expressed genes. *Yn situ* does not produce large, aggregated signals even after overnight reaction. The size of the signal is determined by the structure of the HCR amplification complex formed *in situ*, not by the sequence or the length of the target RNA. Because the HCR reaction is saturated overnight, the size of the *Yn situ* signal is constant across different targets, allowing more quantitative measure at the single molecule level at a larger dynamic range. The expression of odorant receptor gene is expected at ~1-2% of the total mRNA produced. At this level, we still can resolve individual puncta for quantification purpose. In contrast, RNAscope no longer resolved single molecules for highly expressed genes. For 3^rd^ generation HCR to resolve single molecule, 20 or more probe pairs and precise timing of the reaction are required (Choi et al., 2018). In comparison, *Yn situ* can resolve single molecules with as low as three pair of targeting probes. This significantly decreases the cost and investigation time. We have not fully optimized the probe design. It is possible that a single probe pair may produce enough signal for quantification purposes.

Third, the method is simple to perform. Conventional fluorescent *in situ* hybridization requires molecular cloning and the synthesis of long RNA probes. The single stranded RNA probes used for *in situ* hybridizations are prone to degradation by both endogenous RNase from the tissue itself and exogenous RNase from contamination of common reagents used in the lab. *Yn situ* overcomes the problems by using synthetic short DNA oligos, removing the requirement of RNase-free environment. The timing requirement is much relaxed.

Fourth, *Yn situ* only needs a length of 52 nt to detect the target. This length is smaller than most of coding RNA and primary miRNA. This allows the detection of small RNA species, such as microRNA (miRNA), and RNAs that are only targetable by short sequences, such as circular RNA (circRNA). Even though *Yn situ* is not the only method that can detect small RNAs, it is among the very few methods that can detect small RNAs with digital quantification capacity.

Lastly, *Yn situ* offers significant reduction in the cost and time. Only standard desalted oligos are required as primary probes. The cost is less than 1 cent for the primary probes per assay. This method does not require any additional equipment other than the existing molecular cloning and histology devices. A small-scale synthesis of preamplifier is sufficient for hundreds of tests. These factors make the method cost efficient, significantly lower than any commercially available single molecule *in situ* hybridization method. One consideration is the fluorescent metastable hairpin for HCR, which can run up the cost if purchased from the commercial sources. On the hand, the hairpins can be made through well-established amine-NHS ester reactions (Choi et al., 2010). Thus, the cost of this method can be further reduced.

Although *Yn situ* is simple and cheap, it is not merely a poor man’s *in situ* hybridization method. Because of the low cost, robustness, and binary nature of the signals, *Yn situ* has the potentials for further advanced applications such as high throughput automation and multiplexing. The multiplexing includes the simultaneous detection of multiple RNA species and the simultaneous detection of different molecular classes. Simultaneous detection of multiple RNAs can be implemented by the synthesis of additional preamplifiers that do not bind to the same sequence and initiate different pairs of HCR hairpins. 5 orthogonal HCR hairpins have been demonstrated (Choi et al., 2010). In theory, it is possible to design and synthesize at least four more orthogonal preamplifiers. To detect proteins simultaneously with RNAs, the *Yn situ* needs to be performed in prior to the immunohistochemistry. This is because the formamide in the probe hybridization buffer is a denaturant that affects antibody-protein binding. The detection of protein after *in situ* hybridization has been demonstrated previously (Meyer et al., 2017). *Yn situ* protocol does not involve the high annealing temperature of ribonucleic acid probes at 65°C. Low temperature is more feasible for detection of proteins afterwards. Alternatively, the immunostaining of proteins can be performed first with an added step of PFA fixation to crosslink antibodies to the target. It is possible to perform sequential *Yn situ* to increase the detection scale. The realization and scalability of parallelism warrant further investigation.

## Methods

### RESOURCE AVAILABILITY

#### Lead contact

Further information and requests for resources and reagents should be directed to and will be fulfilled by the lead contacts, Yunming Wu and C. Ron Yu.

#### Materials availability

Plasmids generated in this study will be deposited to Addgene for distribution upon publication.

#### Data and code availability

All original data generated in this study will be available for download at Stowers original data repository upon publication. No computer code was used for analysis.

##### EXPERIMENTAL MODEL AND SUBJECT DETAILS

Wildtype CD1 between postnatal day 0 (P0) and P10 pups are used for experiment. Both sexes are randomly assigned to the experiment. All animals were maintained in Stowers LASF with a 14:10 light cycle and provided with food and water ad libitum. Experimental protocols were approved by the Institutional Animal Care and Use Committee at Stowers Institute and in compliance with the NIH Guide for Care and Use of Animals.

##### METHOD DETAILS

###### Oligonucleotides

Oligonucleotides used in this study were listed in the supplementary Table 1. Oligonucleotides used as PCR primers, HCR probes, *Yn situ* blockers, and *Yn situ* probes were purchased from IDT with standard desalting. Fluorescently labeled HCR hairpins were purchased from Molecular Technologies.

###### Synthesis of preamplifier

Plasmid A1-20B1 was synthesized by Thermo Fisher Scientific GeneArt gene synthesis. The plasmid carried kanamycin resistance. The synthesized plasmid was transformed into One Shot TOP10 Electrocomp E. coli (Thermo Fisher Scientific) for further use. Plasmid DNA was purified by miniprep (Zymo Research). Double stranded preamplifier synthesis template was synthesized by PCR using LongAmp PCR polymerase (NEB), non-phosphorylated forward primer, phosphorylated reverse primer. PCR reaction contained 0.05 ng/μL plasmid, 0.5 μM each primer. The PCR program was 95°C for 2 minutes, 37 cycles of 94°C for 30 seconds and 65°C for 1 minute, a final extension at 72°C for 10 minutes. PCR product was purified using DNA spin column (Zymo Research). Single strand preamplifier was synthesized using Guide-it Long ssDNA Production System (Takara Bio) under manufacture’s instruction. Briefly, 5 μg purified PCR products, 5 μL Strandase A Buffer (10X), and 5 μL Strandase Mix A in 50 μL volume was incubated at 37°C for 5 minutes, then 80°C for 5 minutes. The reaction mix was added with 50 μL Strandase B Buffer (10X) and 1μL Strandase Mix B, incubated at 37°C for 5 minutes, 80°C for 5 minutes. The synthesized preamplifier was purified by DNA spin column (Zymo Research).

###### Polymerase chain reaction (PCR)

PCR performed in Figure 1 was conducted under manufacture’s instruction with changes detailed below. For PrimeSTAR, 10 μL PrimeSTAR Max Premix (2X), 0.2 μM of each primer, 1 ng template were used in a 20 μL system. PCR cycles were 35 cycles of 98°C for 10 seconds, 55°C for 15 seconds, and 72°C for 1 minute. For KAPA HiFi, 10 μL of 2X ReadyMix, 0.2 μM of each primer, 1 ng template were used in a 20 μL system. PCR cycles were 1 cycle of 95°C for 3 minutes, 35 cycles of 98°C for 20 seconds, 60°C for 15 seconds, 72°C for 1 minute, 1 cycle of 72°C for 10 minutes. For Q5, 10 μL of 2X master mix, 0.5 μM of each primer, 1 ng template were used in a 20 μL system. PCR cycles were 1 cycle of 98°C for 30 seconds, 35 cycles of 98°C for 10 seconds, 60°C for 30 seconds, 72°C for 1 minute, 1 cycle of 72°C for 10 minutes. For GoTaq, 10 μL of 2X master mix, 0.5 μM of each primer, 1 ng template were used in a 20 μL system. PCR cycles were 1 cycle of 94°C for 2 minutes, 35 cycles of 94°C for 30 seconds, 65°C for 1 minutes, 1 cycle of 72°C for 10 minutes. For LongAmp, different conditions were used. For the experiment in Figure S1A 10 μL of master mix, 0.5 μM of each primer, 1 ng template were used in a 20 μL system. PCR cycles were 1 cycle of 94°C for 30 seconds, 30 cycles of 94°C for 30 seconds, 65°C for 1 minute, 1 cycle of 65°C for 10 minutes. For the experiment in Figure S1B, a gradient from 52°C to 65°C was used for annealing and extension. For the experiment in Figure S1C, two different gradients were used as annealing temperature. One was 52°C to 65°C. The other was 65°C to 72°C. 65°C was used as extension temperature. For the experiment in Figure S1D, 65°C was used for both annealing and extension. Different concentration of primers and templates were used as indicated in the figure. All PCRs were performed in a thermocycler (Biorad).

###### Gel electrophoresis

1% TopVision Agarose gel (Thermo Fisher Scientific) was used for gel electrophoresis analysis DNA molecules were stained with Midori Green Direct DNA staining dye (Bulldog Bio) for gel loading. Electrophoresis was run at 130 V for 30 minutes. Gels were imaged with Gel Logic 100 system (Carestream Health) or SmartDoc gel imaging hood (Stellar Scientific) equipped with an iPhone X (Apple). Images were cropped and contrast enhanced in Fiji.

###### *Yn situ* hybridization

For reproducibility purpose, we attached a step-by-step protocol for performing the procedures.

###### Conventional fluorescent in situ hybridization (FISH)

Conventional fluorescent *in situ* hybridization was conducted following previously described method (Ishii et al., 2004). Briefly, the olfactory epithelia were dissected and embedded in O.C.T. (Sakura Finetek). The embedded samples were snap-frozen in liquid nitrogen. The samples were stored under −70°C until sectioning. The tissue blocks were cut into 10 μm sections using a cryostat (CryoStar NX70) and mounted on charged slides (Thermo Fisher Scientific). The sections were dried on a slide warmer at 100 °C for 2 minutes, fixed with 4% PFA in PBS for 1 hour, fixed with EDC fixative (Pena et al., 2009) for 1 hour before hybridization. Digoxigenin and fluorescein labeled ribonucleotide probes targeting 3’ UTR regions were used. The hybridization was conducted at 65 °C overnight. After washing with SSC, the probes were detected with anti-digoxigenin and anti-fluorescein antibodies conjugated with alkaline phosphatase (AP) and horseradish peroxidase (HRP) using AP detection kit (Roche) and tyramide signal amplification (TSA, Thermo Fisher Scientific) kits. Slides were mounted with No. 1.5 coverslip using Y-mount.

###### RNAscope

RNAscope was performed according to manufacturer’s instruction using probes designed by the company.

###### 3^rd^ generation HCR in situ hybridization

3^rd^ generation HCR was performed according to manufacturer’s instruction using probes designed by the company.

###### Microscopy

Conventional FISH images were taken using Zeiss LSM700 confocal microscope using Plan-Apochromat 20X/0.8 M27 lens. HCR, RNA Scope, and *Yn situ* images were taken using Leica SP8 confocal microscope equipped with HyD hybrid detector using HC PL APO 100X/1.40 Oil lens. Hyvolution images were taken using Leica SP8 confocal microscope under the Hyvolution mode with 0.6 AU pinhole and deconvolved using prolong gold as mounting media in LAS X (Leica). Images were exported as tiff format and analyzed in Fiji. Pixel intensities were measured as procedure generated units from the microscope.

##### QUANTIFICATION AND STATISTICAL ANALYSIS

For SNR calculation, an area in the background was selected to extract pixel intensity values. For signals in the puncta generated by *Yn situ*, RNS-Scope, and HCR, a threshold function in Fiji was used to create masks for the puncta, where the signal intensities for every pixel was extracted. For AP *in situ*, the signals were diffuse. A high signal intensity area was selected without thresholding to extract pixel intensities. The histograms for background and signals were plotted after the signals were normalized to 4096 grayscales. SNR was calculated using the mean values of the signal divided by the variance of the background signal. This calculation avoided the use of background signal intensity because imaging threshold may artificially change the values.

For quantification of signals within a cell, each signal punctum was treated as a single molecule and the number of spots detected in a cell was used to measure the number of RNA molecules in that cell.

The puncta sizes were measured using Hyvolution images. Statistical test in Figure 4C was conducted using ANOVA.

## Supporting information

Yn situ Protocol

## KEY RESOURCES TABLE

**Table 1.**
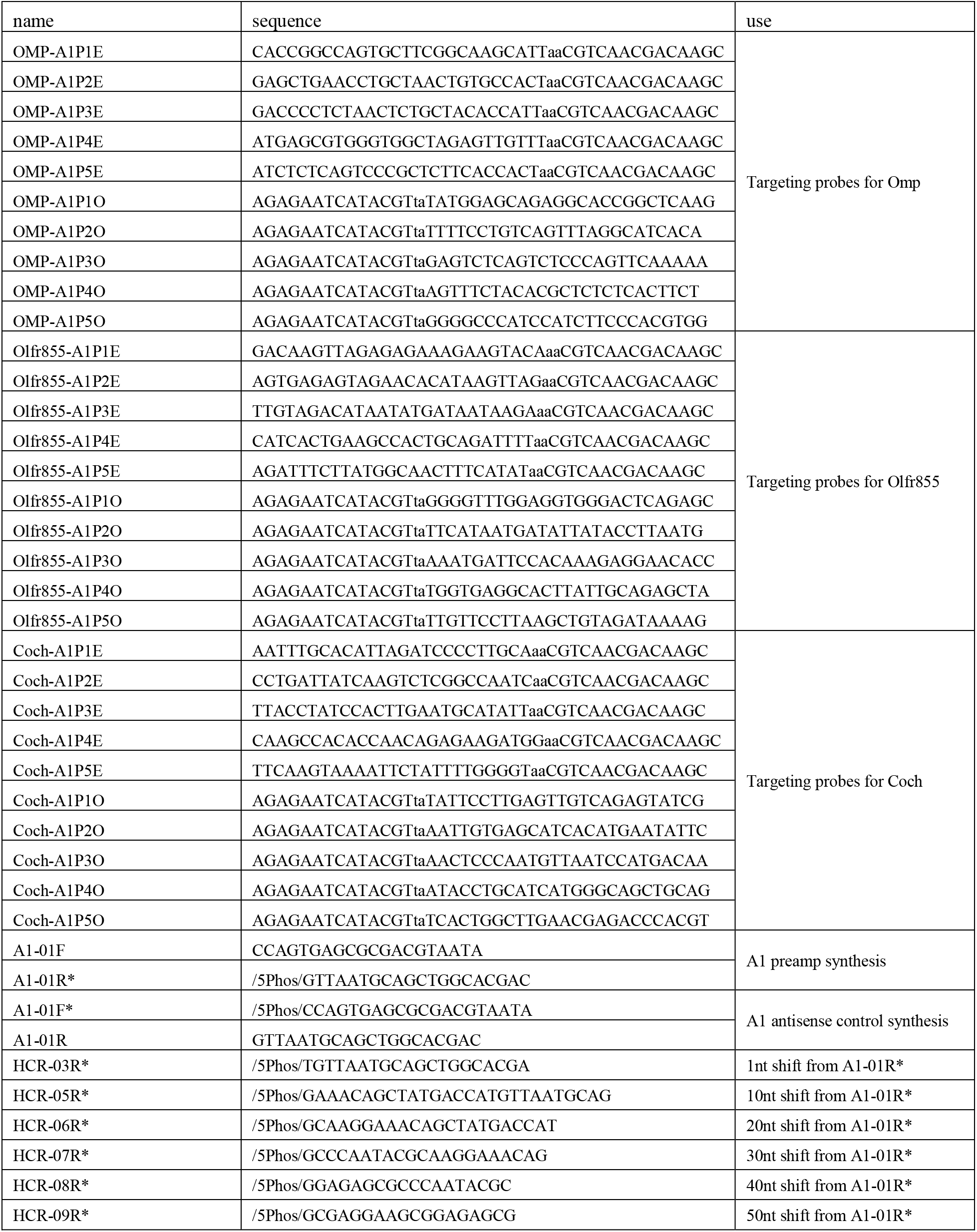
Oligos used in this study

## Supplementary Information

**Figure S1.**
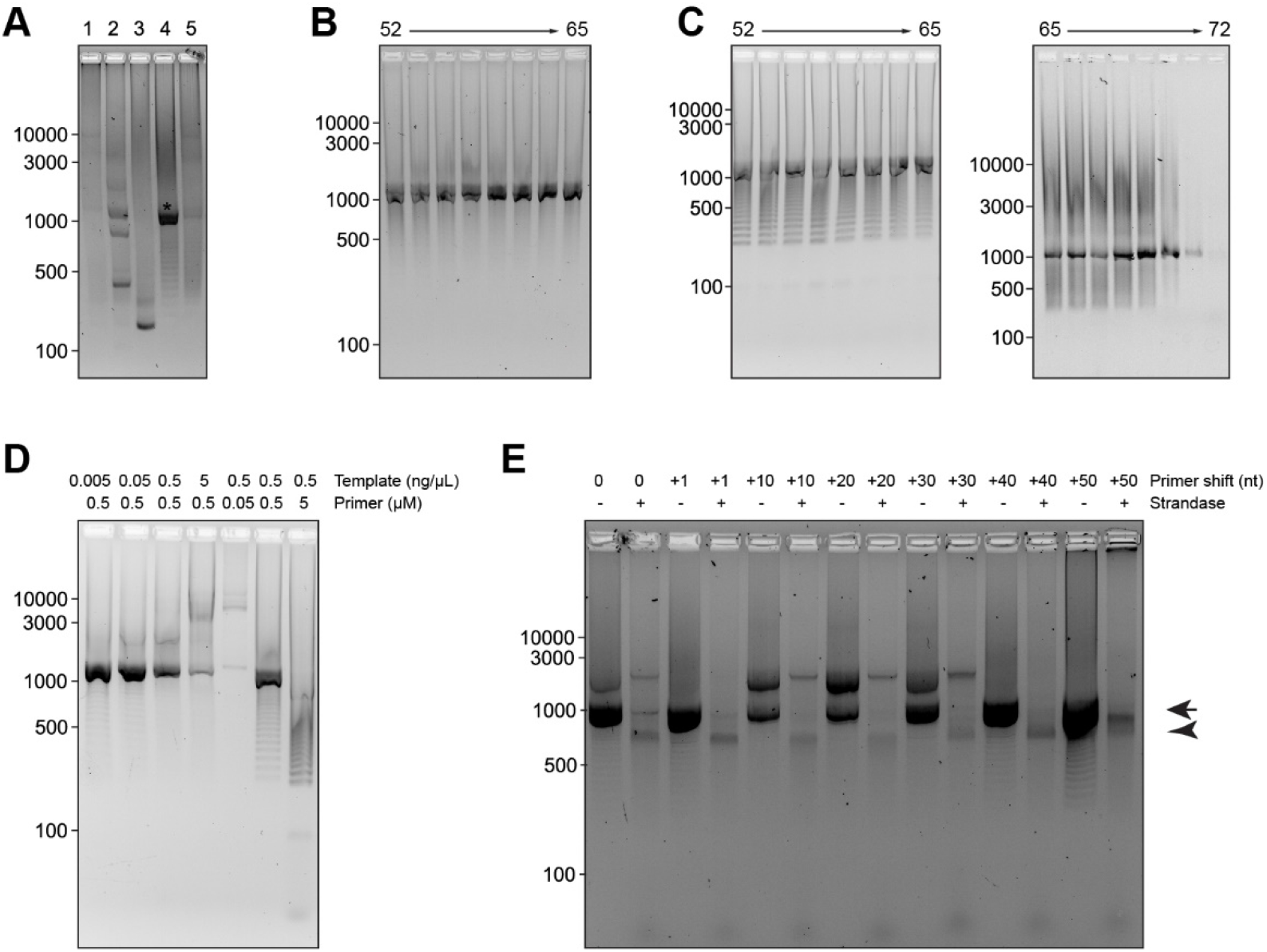
Design and the synthesis of a preamplifier probe. **A**. Performance of different PCR polymerase in the amplification of the pre-amplifier probe sequence. 1. PrimeSTAR HS DNA polymerase. 2. KAPA HiFi DNA polymerase. 3. Q5 DNA polymerase. 4. LongAmp Taq DNA polymerase. 5. GoTaq long PCR master mix. **B.**. Pre-amplifier probe sequence amplification at different temperatures using two-step PCR. **C**. Pre-amplifier probe sequence amplification at different temperatures using three-step PCR. **D**. Pre-amplifier probe sequence amplification with different concentration of template and primers. **E**. Pre-amplifier probe synthesis with different PCR primers. The expected PCR product and ssDNA probe are indicated by arrow and arrowheads, respectively.

