## Supplementary material for "Yn-situ: a robust single RNA molecule *in situ* detection method": Yn situ Protocol

***Yn situ* Hybridization Protocol**

1. Stowers Institute for Medical Research, Kansas City, MO 64110, USA.
2. Department of Anatomy and Cell Biology, University of Kansas Medical Center, 3901 Rainbow Boulevard, Kansas City, KS 66160, USA.
3. Lead Contact

**Probe design**

Probes should be designed against the unique regions of the target RNA. The 52 nt being targeted should be blasted against the transcriptome to determine if there are similar sequences. The GC contents should be within 30-90 %. Repetitive sequences (e.g. AAAAAA and ATATATAT) should avoided.

**Targeting probe preparation**

1. Upon arrive, dissolve individual oligos to 200 µM with TE buffer.
2. Mix all odd probes into one tube to make each probe 2 µM.
3. Mix all even probes into one tube to make each probe 2 µM.
4. Make aliquots and store the aliquots in -20 ℃.

**Tissue Preparation**

1. Sacrifice the animal by cervical dislocation.

2. Dissect the brain and embed the tissue in O.C.T.

3. Float a PCR tube rack on the surface of liquid nitrogen.

4. Place the O.C.T. embedded tissue block on the tube rack.

5. Wait until O.C.T. solidifies.

6. Drop the O.C.T. block into liquid nitrogen until there is no bubble.

7. Store the block in -80 ℃ until use.

**Tissue sectioning**

1. Section the tissue at 10 µm using a cryostat under -25 ℃.

2. Dry slides at 100 ℃ for 2 minutes and proceed directly to fixation.

**Fixation, blocking and permeabilization**

* The steps are conducted at room temperature (RT) if not specified.

1. Fix the slide in 4% PFA for 1 hour.

2. Wash with PBS 3 × 5 minutes.

3. Wash with freshly prepared1-methylimidazole buffer for 5 minutes.

4. Fix the slide in EDC fixative for 1 hour.

5. Wash with PBS for 5 minutes.

6. The slides can be stored after fixation at -80 ℃ for several weeks.

7. Air dry the slides and block the tissue with ImmEdge hydrophobic barrier PAP pen.

8. Digest the tissue with 10 µg/mL proteinase K in TE buffer for 10 min at 40 ℃.

*The optimal proteinase K concentration and digestion time need to be determined for different tissue type and age.

9. Wash with PBS 5 minutes.

**Yn situ hybridization**

1. Wash slides with hybridization solution for 5 minutes.

2. Prepare probe mixture (4 pmol per probe) by adding 2uL of each of the odd and even probes (2 µM stock solution) into 1 mL hybridization buffer.

3. Hybridize at 40 ℃, overnight.

4. Remove probes.

5. Wash in wash buffer, 4 × 30 minutes, 40 ℃.

6. Wash in 5× SSCT, 3 × 5 minutes.

7. Wash in hybridization buffer for 5 minutes.

8. Add 5 µL preamplifier (20 ng/µL) into 500uL probe hybridization buffer (final concentration 0.2 ng/µL).

9. Incubate sections with preamplifier for 5 hours, 40℃.

10. Wash in probe wash buffer for 30 minutes, 40℃.

11. Wash in amplification buffer for 5 minutes.

12. Anneal 10 µL Hairpins (3uM) in a thermocycler, 95 ℃ 90 seconds, -2 ℃/ minutes to 25 ℃.

*The two HCR hairpins need to be annealed separately.

13. Add the 10 µL annealed hairpin into 500 µL amplification buffer (1:50 dilution).

14. Incubate sections in hairpin solutions overnight, RT.

15. Wash in 5× SSCT, 3 × 5 minutes.

16. Mount the slides with coverslips and image the sample under a fluorescent microscope.

**Solution preparation**

*Methylimidazole buffer* (100 mL)

Add 1 mL 1-methlyimidazle to 80 mL H_2_O. Add 12 M HCl to adjust pH to 8.0 (around 200 µL). Add 10mL 3 M NaCl. Add ddH_2_O to 100 mL.

*EDC fixatives*

Add 176 µL EDC to 10 mL imidazole buffer. Add 100 µL 10X 5-ETT. Add 12 M HCl to pH 8.0 (around 100 µL).

*1 M Citric acid*

Dissolve 19.2 g citric acid to 80 mL H_2_O. Add ddH_2_O to 100 mL.

*20X SSC (3 M NaCl, 300 mM Citric acid)*

In 800 ml ddH_2_O (RNase free if required), add 175.3 g NaCl and 88.2 g trisodium citrate. Adjust the pH to 7.0 with 1 M HCl. Adjust the volume to 1 L with ddH_2_O. Sterilize the solution by autoclaving.

*The following buffers, including probe hybridization buffer, amplification buffer, and wash buffer are identical to the ones used in 3^rd^ generation HCR.

*Probe hybridization buffer*

30% formamide, 5× sodium chloride sodium citrate (SSC), 9 mM citric acid (pH 6.0), 0.1% Tween 20, 50 µg/mL heparin, 1X Denhardt’s solution, 10% dextran sulfate

For 40 mL of solution

12 mL deionized formamide

10 mL of 20× SSC

360 µL 1 M citric acid, pH 6.0

400 µL of 10% Tween 20

200 µL of 10 mg/mL heparin

800 µL of 50× Denhardt’s solution

8 mL of 50% dextran sulfate

Fill up to 40 mL with ddH_2_O.

*Probe wash buffer*

30% formamide, 5× SSC, 9 mM citric acid (pH 6.0), 0.1% Tween 20, 50 µg/mL heparin

For 1L of solution

300 mL deionized formamide

250 mL of 20× SSC

9 mL 1 M citric acid, pH 6.0

10 mL of 10% Tween 20

5 mL of 10 mg/mL heparin

Fill up to 1 L with ddH_2_O

*100X 5-ETT*

Dissolve 2 g 5-ETT in 1.5 mL ddH_2_O

*Amplification buffer*

5× SSC, 0.1% Tween 20, 10% dextran sulfate

For 40 mL of solution

10 mL of 20× SSC

400 µL of 10% Tween 20

8 mL of 50% dextran sulfate

Fill up to 40 mL with ddH_2_O
